## Supplemental Figures for "Clusters of deep intronic RbFox motifs embedded in large assembly of splicing regulators sequences regulate alternative splicing"

Suppl. Fig. 1

| Chromosome | Start | End | -log10(P-value) | Log2 Fold Change | Strand | Gene | Ensembl ID | Feature |
|---|---|---|---|---|---|---|---|---|
| chrTrkB_Ntrk2_minigene | 31,100 | 31,115 | 400 | 3.14437648711126 | + | Ntrk2 | ENSMUSG00000055254 | Distal intron |
| chrTrkB_Ntrk2_minigene | 45,552 | 45,584 | 400 | 3.20890405198609 | + | Ntrk2 | ENSMUSG00000055254 | Distal intron |
| chrTrkB_Ntrk2_minigene | 76,182 | 76,233 | 38.9618171340093 | 3.77851130406743 | + | Ntrk2 | ENSMUSG00000055254 | Distal intron |
| chrTrkB_Ntrk2_minigene | 122,670 | 122,695 | 12.4330678000343 | 3.89923278816692 | + | Ntrk2 | ENSMUSG00000055254 | Distal intron |
| chrTrkB_Ntrk2_minigene | 142,283 | 142,334 | 21.888569448234 | 4.13082488341521 | + | Ntrk2 | ENSMUSG00000055254 | Distal intron |
| chrTrkB_Ntrk2_minigene | 142,334 | 142,378 | 15.5991167844516 | 3.89343880102932 | + | Ntrk2 | ENSMUSG00000055254 | Distal intron |
| chrTrkB_Ntrk2_minigene | 146,090 | 146,137 | 23.8935711052066 | 3.69845068266664 | + | Ntrk2 | ENSMUSG00000055254 | Distal intron |

**Suppl. Fig. 2**

**A**

| Rank | Motif | P-value | log P-pvalue | % of Targets | % of Background | STD(Bg STD) |
|---|---|---|---|---|---|---|
| 1 | 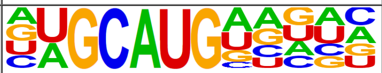 | 1e-2765 | -6.368e+03 | 32.51% | 7.72% | 30.4bp (34.4bp) |
| 2 | 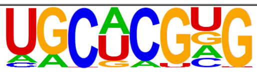 | 1e-155 | -3.570e+02 | 1.41% | 0.22% | 25.4bp (15.7bp) |
| 3 | 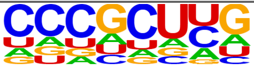 | 1e-146 | -3.372e+02 | 6.32% | 3.10% | 35.7bp (30.9bp) |
| 4 | 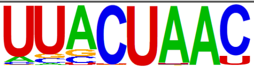 | 1e-132 | -3.048e+02 | 1.41% | 0.27% | 32.9bp (24.5bp) |
| 5 | 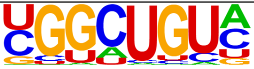 | 1e-127 | -2.945e+02 | 15.36% | 10.40% | 33.1bp (37.8bp) |

**B**

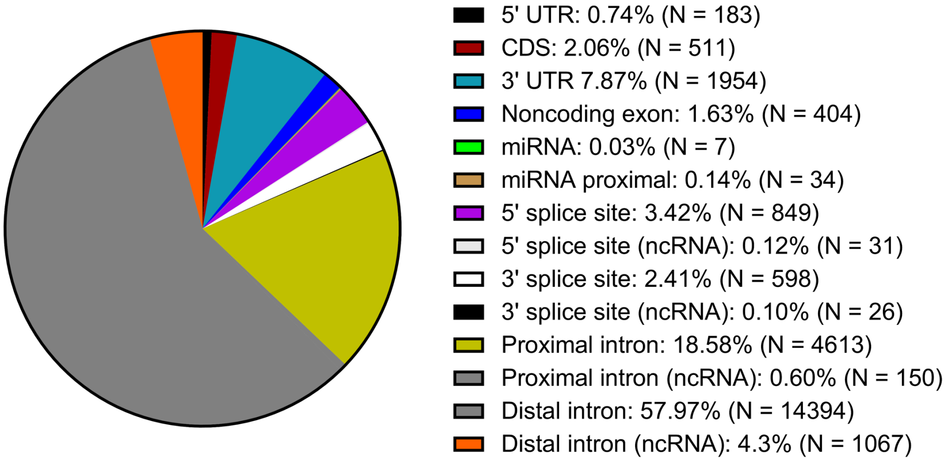


**C**

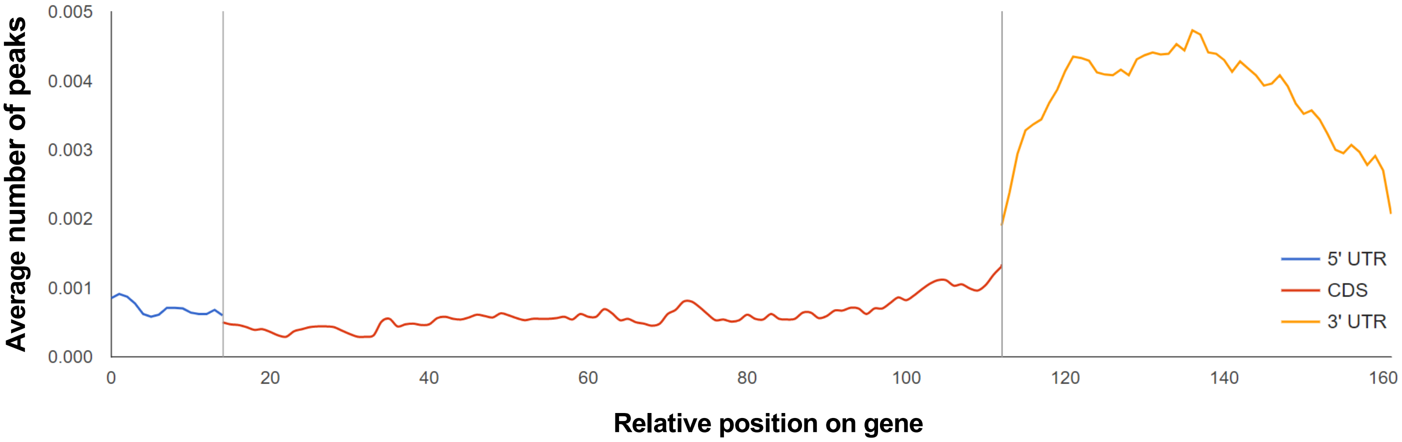


**D**

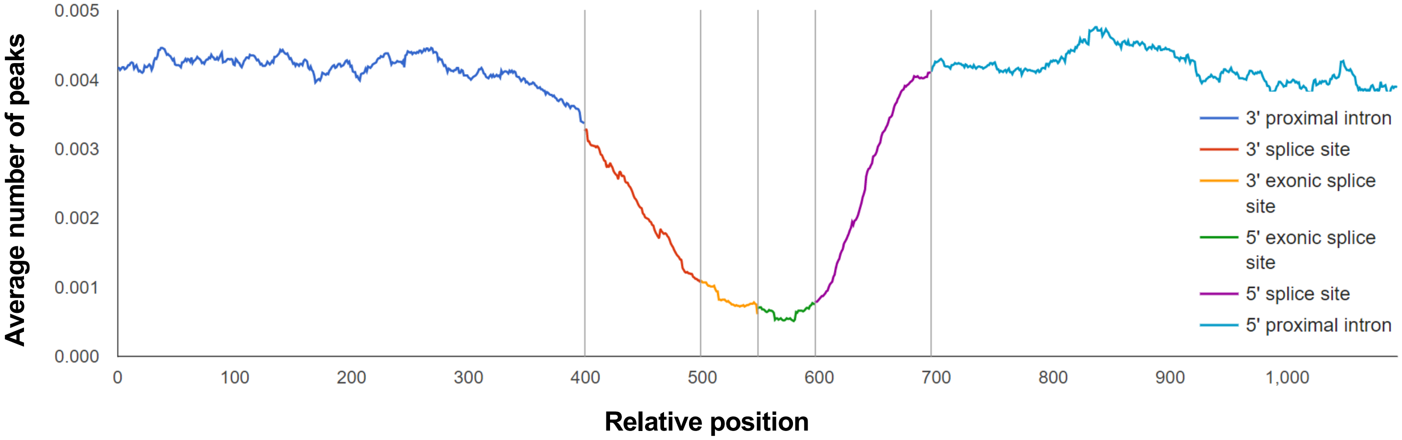

### Suppl. Fig. 3

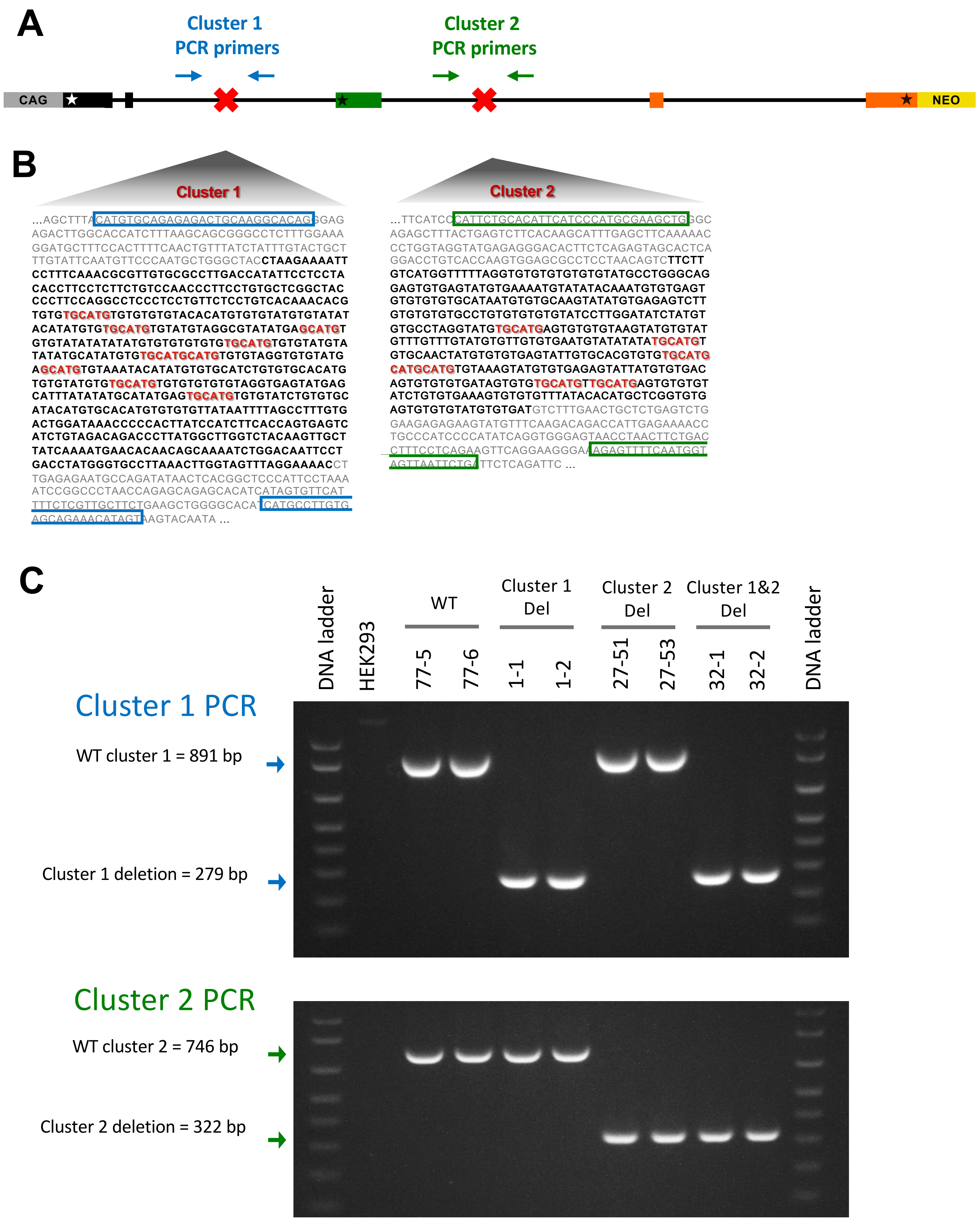

A
Cluster 1
PCR primers
Cluster 2
PCR primers
CAG
NEO
B
Cluster 1
...AGCTTTACATGTGCAGAGAGACTGCAAGGCACAGGGAG
AGACTTGGCACCATCTTTAAGCAGCGGGCCTCTTTGGAAA
GGATGCTTTCCACTTTTCAACTGTTTATCTATTTGTACTGCT
TTGTATTCAATGTTCCCAATGCTGGGCTACCTAAGAAAATT
CCTTTCAAACGCGTTGTGCGCCTTGACCATATTCCTCCTA
CACCTTCCTCTTCTGTCCAACCCTTCCTGTGCTCGGCTAC
CCCTTCCAGGCCTCCCTCCTGTTCTCCTGTCACAAACACG
TGTGTGCATGTGTGTGTGTACACATGTGTGTATGTGTATAT
ACATATGTGTGCATGTGTATGTAGGCGTATATGAGCATGT
GTGTATATATATATGTGTGTGTGTGTGCATGTGTGTATGTA
TATATGCATATGTGTGCATGCATGTGTGTAGGTGTGTATG
AGCATGTGTAAATACATATGTGTGCATCTGTGTGCACATG
TGTGTATGTGTGCATGTGTGTGTGTGTAGGTGAGTATGAG
CATTTATATATGCATATGAGTGCATGTGTGTATCTGTGTGC
ATACATGTGCACATGTGTGTGTTATAATTTTAGCCTTTGTG
ACTGGATAAACCCCCACTTATCCATCTTCACCAGTGAGTC
ATCTGTAGACAGACCCTTATGGCTTGGTCTACAAGTTGCT
TATCAAAATGAACACAACAGCAAAATCTGGACAATTCCT
GACCTATGGGTGCCTTAAACTTGGTAGTTTAGGAAAACCT
TGAGAGAATGCCAGATATAACTCACGGCTCCCATTCCTAAA
ATCCGGCCCTAACCAGAGCAGAGCACATCATAGTGTTCAT
TTTCTCGTTGCTTCTGAAGCTGGGGCACATCATGCCTTGTG
AGCAGAAACATAGTAAGTACAATA ...
Cluster 2
...TTCATCCCATTCTGCACATTCATCCCATGCGAAGCTGGGC
AGAGCTTTACTGAGTCTTCACAAGCATTTGAGCTTCAAAAAC
CCTGGTAGGTATGAGAGGGACACTTCTCAGAGTAGCACTCA
GGACCTGTCACCAAGTGGAGCGCCTCCTAACAGTCTTCTT
GTCATGGTTTTTAGGTGTGTGTGTGTATGCCTGGGCAG
GAGTGTGAGTATGTGAAAATGTATATACAAATGTGTGAGT
GTGTGTGTGCATAATGTGTGCAAGTATATGTGAGAGTCTT
GTGTGTGTGCCTGTGTGTGTGTATCCTTGGATATCTATGT
GTGCCTAGGTATGTGCATGAGTGTGTGTAAGTATGTGTAT
GTTTGTTTGTATGTGTTGTGTGAATGTATATATATGCATGT
GTGCAACTATGTGTGTGAGTATTGTGCACGTGTGTGCATG
CATGCATGTGTAAAGTATGTGTGAGAGTATTATGTGTGAC
AGTGTGTGTGATAGTGTGTGCATGTTGCATGAGTGTGTGT
ATCTGTGTGAAAGTGTGTGTTTATACACATGCTCGGTGTG
AGTGTGTGTATGTGTGATGTCTTTGAACTGCTCTGAGTCTG
GAAGAGAGAAGTATGTTTCAAGACAGACCATTGAGAAAACC
TGCCCATCCCCATATCAGGTGGGAGTAACCTAACTTCTGAC
CTTTCCTCAGAAGTTCAGGAAGGGAAAGAGTTTTCAATGGT
AGTTAATTCTGATTCTCAGATTC ...
C
DNA ladder
HEK293
WT
77-5
77-6
Cluster 1
Del
1-1
1-2
Cluster 2
Del
27-51
27-53
Cluster 1&2
Del
32-1
32-2
DNA ladder
Cluster 1 PCR
WT cluster 1 = 891 bp
Cluster 1 deletion = 279 bp
Cluster 2 PCR
WT cluster 2 = 746 bp
Cluster 2 deletion = 322 bp

**Suppl. Fig. 4**

**A**

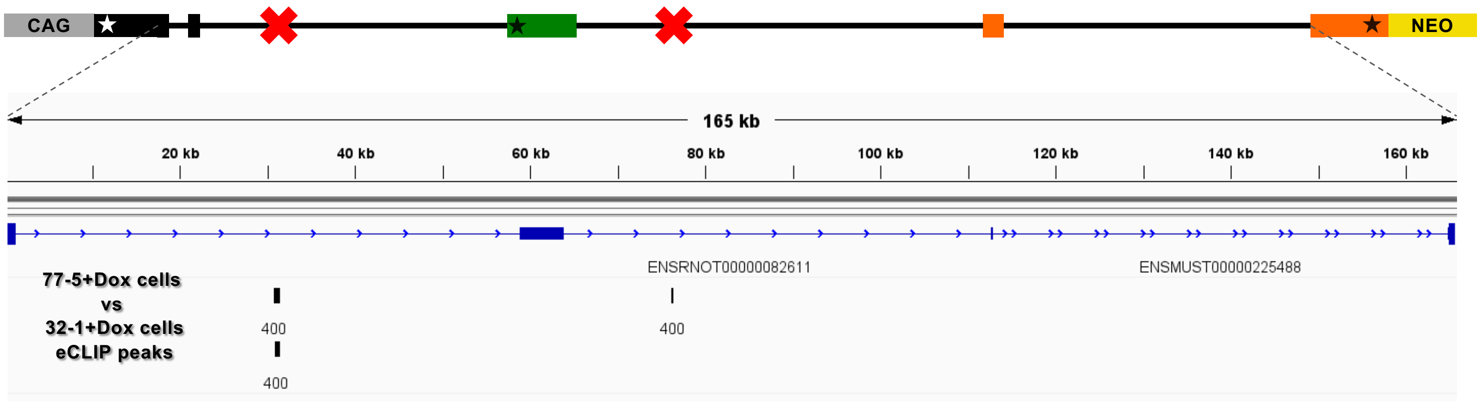


**B**

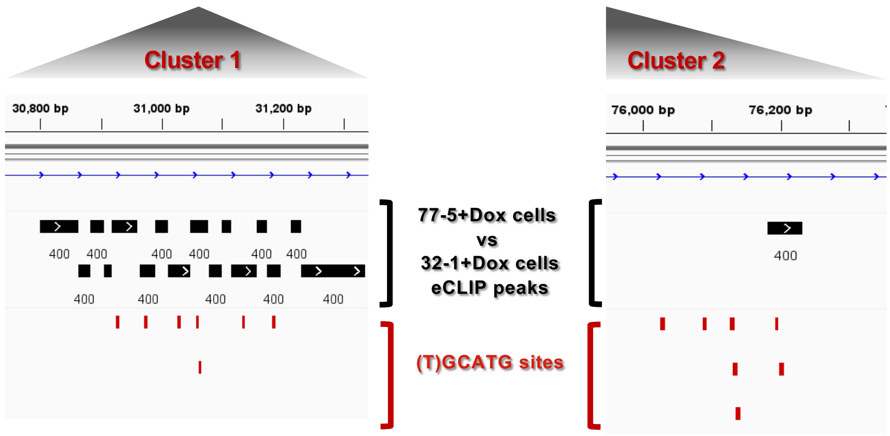


**C**

| Chromosome | Start | End | -log10(P-value) | Log2 Fold Change | Strand | Gene | Ensembl ID | Feature |
|---|---|---|---|---|---|---|---|---|
| chrTrkB_Ntrk2_minigene | 30,801 | 30,863 | 400 | 7.2027601433326 | + | Ntrk2 | ENSMUSG00000055254 | Distal intron |
| chrTrkB_Ntrk2_minigene | 30,863 | 30,882 | 400 | 7.83245626508985 | + | Ntrk2 | ENSMUSG00000055254 | Distal intron |
| chrTrkB_Ntrk2_minigene | 30,882 | 30,906 | 400 | 8.29713507889578 | + | Ntrk2 | ENSMUSG00000055254 | Distal intron |
| chrTrkB_Ntrk2_minigene | 30,906 | 30,918 | 400 | 7.94503411560365 | + | Ntrk2 | ENSMUSG00000055254 | Distal intron |
| chrTrkB_Ntrk2_minigene | 30,918 | 30,961 | 400 | 5.09097712399694 | + | Ntrk2 | ENSMUSG00000055254 | Distal intron |
| chrTrkB_Ntrk2_minigene | 30,964 | 30,990 | 400 | 5.11980751567679 | + | Ntrk2 | ENSMUSG00000055254 | Distal intron |
| chrTrkB_Ntrk2_minigene | 30,990 | 31,011 | 400 | 4.72464717782695 | + | Ntrk2 | ENSMUSG00000055254 | Distal intron |
| chrTrkB_Ntrk2_minigene | 31,011 | 31,047 | 400 | 3.69840522735328 | + | Ntrk2 | ENSMUSG00000055254 | Distal intron |
| chrTrkB_Ntrk2_minigene | 31,047 | 31,076 | 400 | 4.83377754237091 | + | Ntrk2 | ENSMUSG00000055254 | Distal intron |
| chrTrkB_Ntrk2_minigene | 31,078 | 31,100 | 400 | 6.95832944582644 | + | Ntrk2 | ENSMUSG00000055254 | Distal intron |
| chrTrkB_Ntrk2_minigene | 31,100 | 31,115 | 400 | 5.96082696528812 | + | Ntrk2 | ENSMUSG00000055254 | Distal intron |
| chrTrkB_Ntrk2_minigene | 31,115 | 31,156 | 400 | 4.41839823542178 | + | Ntrk2 | ENSMUSG00000055254 | Distal intron |
| chrTrkB_Ntrk2_minigene | 31,156 | 31,174 | 400 | 7.31839724537609 | + | Ntrk2 | ENSMUSG00000055254 | Distal intron |
| chrTrkB_Ntrk2_minigene | 31,174 | 31,197 | 400 | 6.44863628125885 | + | Ntrk2 | ENSMUSG00000055254 | Distal intron |
| chrTrkB_Ntrk2_minigene | 31,214 | 31,231 | 400 | 6.04066610949988 | + | Ntrk2 | ENSMUSG00000055254 | Distal intron |
| chrTrkB_Ntrk2_minigene | 31,231 | 31,337 | 400 | 8.14300321860379 | + | Ntrk2 | ENSMUSG00000055254 | Distal intron |
| chrTrkB_Ntrk2_minigene | 76,182 | 76,233 | 400 | 5.08149532441559 | + | Ntrk2 | ENSMUSG00000055254 | Distal intron |

Suppl. Fig. 5

A

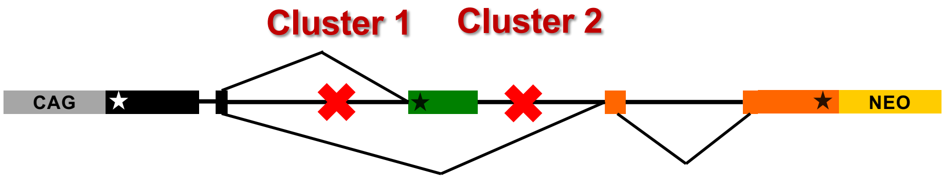


B

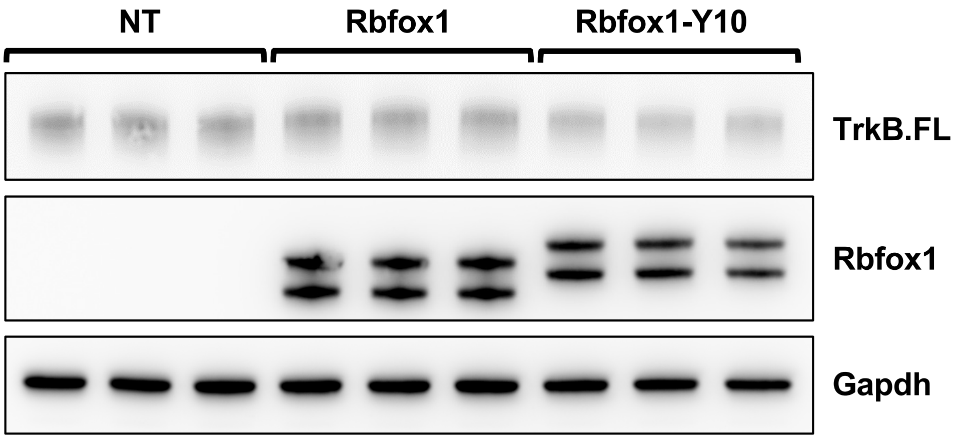
